# Treatment of age-related decreases in GTP levels restores endocytosis and autophagy

**DOI:** 10.1101/2025.06.09.658689

**Authors:** R. A. Santana, J. M. McWhirt, G. J. Brewer

## Abstract

Age-related declines in neuronal bioenergetic levels may limit vesicular trafficking and autophagic clearance of damaged organelles and proteins. Age-related ATP depletion would impact cognition dependent on ionic homeostasis, but limits on proteostasis powered by GTP are less clear. We used neurons isolated from aged 3xTg-AD Alzheimer’s model mice and a novel genetically encoded fluorescent GTP sensor (GEVAL) to evaluate live GTP levels in situ. We report an age-dependent reduction in ratiometric measurements of free/bound GTP levels in living hippocampal neurons. Free-GTP co-localized in the mitochondria decreased with age accompanied by the accumulation of free-GTP labeled vesicular structures. The energy dependence of autophagy was demonstrated by depletion of GTP with rapamycin stimulation, while bafilomycin inhibition of autophagy raised GTP levels. 24 hr. supplementation of aged neurons with the NAD precursor nicotinamide and the Nrf2 redox modulator EGCG restored GTP levels to youthful levels and mobilized endocytosis and lysosomal consumption for autophagy via the respective GTPases Rab7 and Arl8b. This vesicular mobilization promoted the clearance of intraneuronal Aβ aggregates and lowered protein oxidative nitration in AD model neurons. Our results reveal age– and AD-related neuronal GTP energy deficits that impair autophagy and endocytosis. GTP deficits were remediated by an external NAD precursor together with a Nrf2 redox modulator which suggests a translational path.

## INTRODUCTION

Age is the largest risk factor for Alzheimer’s disease (AD), but the age-related mechanisms that increase risk for AD are unclear. Although a proposed lifelong buildup of extracellular amyloid or ROS damage could eventually reach a clinical threshold for dementia, neither mouse models nor human anti-amyloid or antioxidant therapies have validated this proposition. The amyloid-precursor protein is an integral membrane protein that is processed into the aggregation-prone Aβ peptide, beginning in the plasma membrane and continuing in endocytosis [1]; however, little is known about the age-related availability of energy to power endocytosis. We propose a mechanism to better explain the downstream pathophysiological events in AD powered by age-related intracellular aggregation of amyloid-β (Aβ) in the vesicular trafficking system initiated by an oxidative shift in redox state that lowers bioenergetic capacity [2, 3].

Aging and age-related disease are tightly linked to a decline in energetic capacity, a supply and demand imbalance [4]. In the aging brain, ATP levels decline with age, which impacts cognition [5]. Further, high energetic demand in the hippocampus and entorhinal cortex for memory and learning plasticity likely generates high levels of ROS [6] and the need for robust autophagic control of proteostasis, largely powered by GTP [7]. Among nucleotide tri-phosphates like ATP, GTP hydrolysis selectively drives energy-dependent cellular processes of protein synthesis and vesicular trafficking, including endocytosis and autophagy [7]. Levels of GTP depend on the redox state of the electron acceptors and donors NAD^+^/NADH and their concentrations. GTP-driven endocytosis involves the GTPases Rab5 in early endosomes, and Rab7 in late endosomes that fuse with lysosomes using Arl8 GTPase for autophagic clearance. Impairments in the supply of GTP could influence the age-related net production and clearance of amyloid (Aβ42 and Aβ45) resulting in accumulation of early endosomes containing Rab5 on their surface, Rab7decorated late endosomes, and p62 associated autophagosomes as we have seen in neurons from old AD mice [1]. Impairment of the vesicular process of autophagy in AD [8] could be caused by an age-related decline in GTP substrate to the GTPases that control each stage of autophagy [7]. To establish this hypothesis, we evaluated the effects of an energy-boosting small molecule together with an inducer of a balanced redox response.

The redox energy supply of nicotinamide adenine dinucleotide (NAD^+^) is vital to glycolysis and the tricarboxylic acid cycle (TCA), forming free NADH (reduced form) in the mitochondria. NADH is the primary electron donor for oxidative phosphorylation in the brain, for ATP production, but also some ROS. The NAD^+^/NADH redox pair modulates gene expression through redox-sensitive transcription factors [9], acts as a precursor of multiple NAD^+^-dependent enzymes involved in synaptic plasticity and neuronal stress resistance, and regulates crucial signaling pathways related to neuronal survival [10]. A decrease in NAD^+^ synthesis accompanied by high NAD^+^ consumption with age that leads to a reduction of the NAD pool [11].

Nicotinamide is a precursor to NAD^+^ through NAMPT and NMNAT enzymes [10]. NAD^+^ is rapidly converted to NADH by the malate shuttle and mitochondrial transhydrogenase to establish redox balance and power oxidative phosphorylation. A redox shift is a change in the bioenergetic supply of electrons, leading to changes in the intracellular ratio of NAD^+^/NADH as redox currency, oxidized glutathione over GSH (GSSG/GSH) as redox buffer, and plasma cystine/cysteine (cySS/cys) as systemic buffer.

Therefore, a boost with nicotinamide may improve the age-related energetic deficits associated with redox imbalance. We propose that an age-related oxidative redox shift overwhelms energetic capacity to promote pathogenic impairment of proteostasis and downstream synapse loss and cognitive decline in AD. Further, we test whether this decline in proteostasis and Aβ processing can be restored in primary neurons from aged mice by treatment with nicotinamide to boost NAD^+^ levels and with EGCG to engage Nrf2 signaling and redox balance.

Aging and AD are also associated with increased oxyradical damage to proteins, lipids and nucleic acids [12,13]. Boosting energy supplies produces excess free radicals from increased oxidative phosphorylation [14] which may not be neutralized if the cells have been operating in a low redox capacity with low activation of Antioxidant Response Elements (ARE) genes. As transcription factor activator of ARE, Nrf2 resides in the cytoplasm in an inactive state bound to KEAP [4,15]. Exposure to electrophiles releases Nrf2 from KEAP to allow translocation to the nucleus, and activation of ARE-dependent genes such as NAD(P)H quinone oxidoreductase (NQO1), glutathione transferase (GST), thioredoxin reductase (TRX), gamma-glutamyl cysteine synthase, malic enzyme and catalase [16]. EGCG, the primary polyphenol in green tea, is a safe and widely studied Nrf2 inducer [17]. Therefore, we compared the effect on GTP levels of EGCG as a Nrf2 inducer plus nicotinamide for naturally regulating ROS levels while boosting NAD^+^ energy to improve endocytosis and autophagy in primary hippocampal neurons across the age-span from non-transgenic control and 3xTg-AD mice. This approach represents an opportunity to decelerate, stop, or reverse aging and AD by therapeutic interventions with these small molecules. We introduced these concepts in a perspective review [7].

## Material and Methods

### Cell culture

Primary neurons were obtained from the triple transgenic mouse model of AD (3xTg-AD) with human βAPP (SWE), PS1 (M146V), and Tau (P301L) transgenes (Oddo [18] 2003) in three different age ranges: young (2 – 6 months), middle (8 – 11 months) and old (17 – 28 months), around the median lifespan. Non-transgenic mice (NTg; C57/Bl6, Charles River with wild-type transhydrogenase) of the same age ranges were used as controls. All mice were genotyped from tail snips before use (Transnetyx, Cordova, TN). All animal procedures were approved by the Institutional Animal Care and Use Committee (AUP-23-13) and performed according to regulations. Culture of isolated adult neurons was as previously described [19]. Briefly, isoflurane by inhalation was used to anesthetize the mice. From the brains, the hippocampi with entorhinal cortices were combined from both hemispheres and sliced at 0.5 mm in Hibernate AB (Transnetyx/BrainBits, #HAB 500, osmolarity adjusted to 298 mOsm with 5 M NaCl). Slices were placed in a 30°C bath for 8 min. The tissue was digested with 2 mg/ml papain (Transnetyx/BrainBits, #52D234) in Hibernate A minus calcium (TransnetyxBrainBits HACA) and 0.5 mM Glutamax (Thermo Fisher/GIBCO, Grand Island, NY) for 30 min at 30°C in a shaking water bath (170 rpm). The papain solution was replaced with Hibernate AB before trituration to isolate, the cells from each hippocampus and entorhinal cortex. The cell suspension was transferred to the top of a 15 mL tube prepared with four layered densities of Optiprep (Cosmo Bio, Carlsbad, CA, AXS-1114542). The gradient was centrifuged at 800 g for 15 min. The neuron-enriched fractions were collected and transferred to 5 mL of Hibernate AB. The cell suspension was centrifuged for 3 min at 300 g and the supernatant discarded. After resuspension of the cell pellet in culture medium, the cells were plated at 32,000 cells/cm^2^ onto 15 mm coverslips (Carolina Biologicals, Burlington, NC, 41001112), precoated with poly-D-lysine (Sigma Aldrich, St Louis, MO, P6407-5MG). Culture medium was Neurobasal A with B27 (Thermo Fisher/GIBCO (10888022, 17-504-044), 1 mM Glutamax, supplemented with 5 ng/mL each of mouse FGF2 (Pepro-Tech 450-33) and PDGFBB (Pepro-Tech 315-18) for trophic support. The medium was adjusted to 290 mOsm with 5 M NaCl. One-half medium changes were made on days 3, and 6 with 10 ng/mL growth factors, assuming consumption of the prior growth factors, and on day 10 without growth factors. The cells were cultured for 12–15 days at 37° C in 5% CO_2_ and 9% O_2_ at saturated humidity (Thermo-Forma, Marietta, OH, Model 3130).

### GTP ratio measurements

GTP evaluator (GEVAL) plasmids were a gift from Dr. Mikhail A. Nikiforov (Duke University, Durham, NC) [20]. GEVAL530, a specific GTP-binding construct, has a bacterial FeoB G-protein fused to a YFP sensor with a K_eff_=528 µM to cover the cellular range of GTP from 300-800 µM. Plasmids were expanded and purified using the QIAprep Spin Miniprep Kit (QIAGEN, # 27106) according to the manufactureŕs protocol. GEVALnull was used as an unresponsive negative control.

### Neuronal transfection

Hippocampal neurons were transfected with GEVAL plasmids using jetOPTIMUS transfection reagent according to the manufactureŕs specifications (PolyPlus, CA). jetOPTIMUS proved more efficient than Lipofectamine 2000, LipoJet, and Lipo293). Briefly, 0.5 µg plasmid DNA was diluted into 100 µl of jetOPTIMUS buffer and gently mixed (pipette up and down five times). The mixture was added to 0.5 µl of jetOPTIMUS reagent at room temperature onto the DNA solution (ratio 1:1 corresponding to µg DNA: µl reagent), gently mixed, and incubated for 10 min at 22° C. After removing half of the culture medium, neurons at 60-70% confluency were transfected by adding 100 µl transfection mix per 1.76 cm^2^ well dropwise onto the cells. For uniform distribution, the plate was gently shaken back and forth and from side to side. After incubating the cells at 37°C in 5% CO_2_, 9% O_2_ for 4 hr., we added an additional 0.5 ml of pre-warmed complete culture medium. GEVAL signal was analyzed after 24 hr. by confocal imaging.

### Treatments

After 12-14 days in culture, hippocampal neurons were treated for 16 hr. with 2 mM Nicotinamide (Sigma-Aldrich, # N7004; from 200 mM stock in Hibernate A) and 2 µM EGCG (Sigma-Aldrich, # E4143; from 200 µM stock in sterile 18-Mohm water) or combination. Operationally, 200 µl conditioned medium from a 500 µl culture well was transferred to a 0.6 ml sterile tube; drug was added, mixed and returned to the well.

### Cell imaging and analysis

30 min before imaging, cells were rinsed with 37°C ERB buffer (120 mM NaCl, 3.5 mM KCl, 1.2 mM CaCl_2_, 1 mM MgCl_2_, 0.4 mM KH_2_PO_4_, 5 mM NaHCO_3_, 1.2 mM NaSO_4_, 20 mM HEPES, 5 mM Glucose, pH 7.4 and adjusted at 290 mOsm). Cells were imaged with an LFD-LSM880 Zeiss microscope equipped with a multi-line Argon laser, and 405 nm diode laser with plan-Apochromat 63x/NA 1.4 oil DIC M27 immersion lens. In each of 10 fields, GTP sensors were excited sequentially with the 405 nm wavelength of the diode laser to measure bound GTP (405-430 nm, laser power at 2.0), and the 488 nm line of the Argon laser to measure unbound (free) GTP (laser power at 6.0). Fluorescent emission was collected from 493 to 554 nm (Fig. 1A) (512 x 512 pixels and 67.48 x 67.48 µm). Sequential imaging from the same fields was carried out over 25 min. in experiments with bafilomycin and rapamycin. Image analysis was performed using ImageJ/Fiji (v1.53t, NIH). Background was corrected using the rolling ball feature at a size of 10 pixels before calculating ratios. According to Bianchi-Smiraglia & Nikiforov [21], pixel-by-pixel ratiometric images were created to analyze GTP distribution within cells and One brightness setting of +30% was used for all figure images. However, for quantitative image analysis, all images within the same experiment were analyzed at their original intensities at constant threshold. Images of the same field at the two excitations were aligned as a stack using the Stack Reg plug-in. In the Plugins menu, the Ratio Plus plug-in was selected, using ex488 as Image 1 and ex405 as Image 2, and a Multiplication factor of 1 with enhance contrast at 0.4%. For display (not quantitative analysis), the false-color image used the Fire option in the LookupTables (LUT) menu at common brightness and contrast settings.

**Fig. 1.**
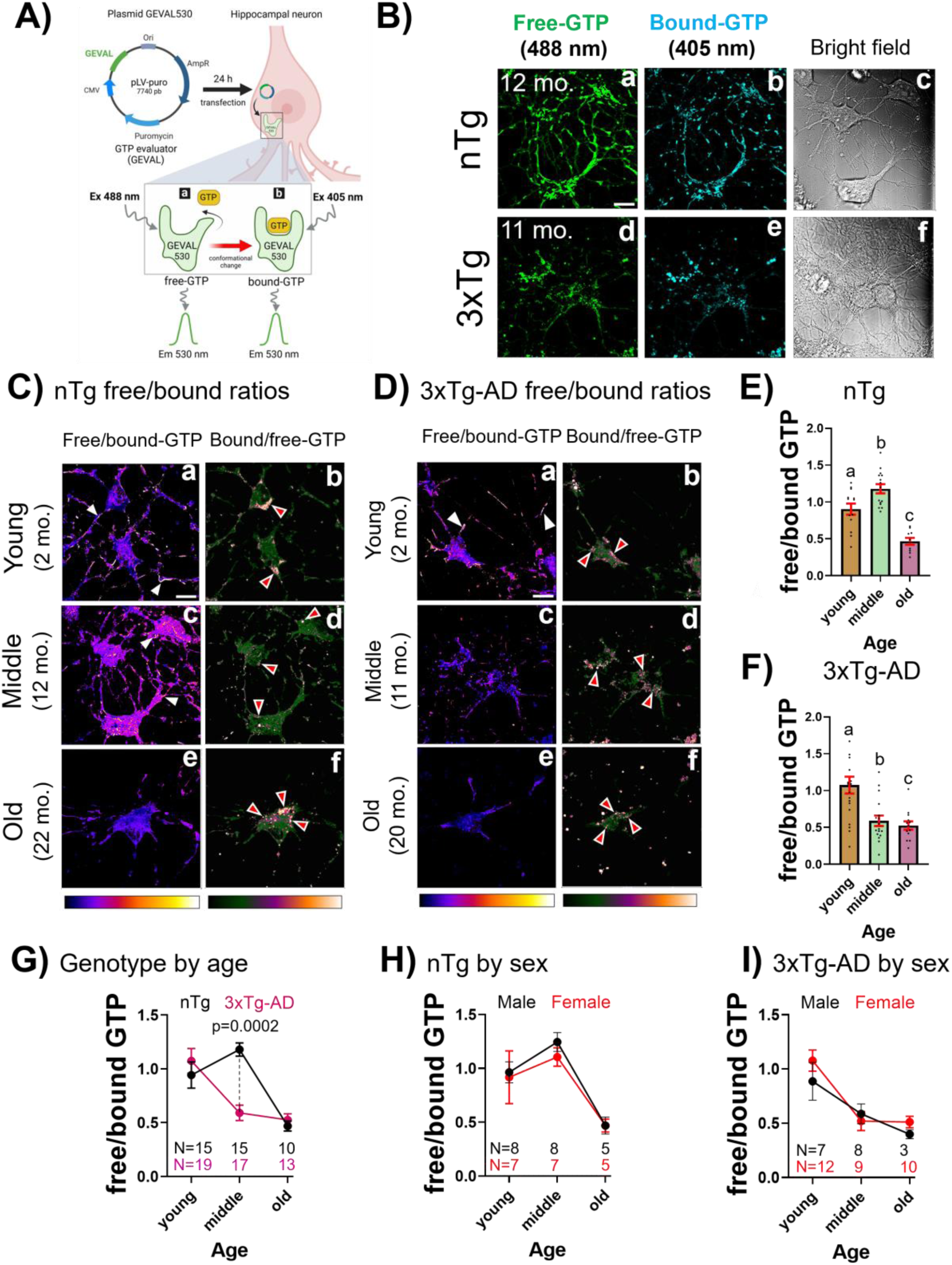
Aging and AD-genotype-related decreases in free GTP in hippocampal neurons. A) Hippocampal neurons transfected with GEVAL530 plasmid for 24 h. GTP sensor undergoes a GTP-driven conformational change upon binding to GTP. Transfected cells were excited sequentially with a) 488 nm to measure free (unbound) GTP and b) 405 nm to measure bound GTP from fluorescent emission at 530 nm. Image created with BioRender. B) Representative images of free-GTP after excitation at 488 nm and bound-GTP after excitation at 405 nm in neurons from middle age mice of both genotypes (12 months for nTg (a-c) and 11 months for 3xTg-AD (d-f)). Bright field shows the axonal-dendritic network in the neurons (c,f). Scale bar 10 μm. C) Representative images of the ratio of free/bound and bound/free from nTg neurons from young (a,b); middle age (c,d) and old age mice (e-f). D) Similar images of 3xTg-AD neurons from young (a,b), middle age (c,d) and old age mice (e,f) (20 months). False-colored images are pixel by pixel ratios of hippocampal neurons using Look-Up Table Fire for free/bound and blue-orange-icb for bound/free GTP. Scale bar 10 μm. White arrow heads indicate high free-GTP in axons, dendrites and in somata. Red arrow heads indicate bound-GTP localization at periphery of neuron somata. E) In nTg neurons, middle-age increase in free/bound GTP levels and decrease in in old-age. F) In 3xTg-AD neurons, free/bound GTP levels decline in middle age with respect to young age and remain low in old age. In E, F, different letters indicate subgroup differences at p<0.0001 after Tukey correction for multiple comparisons. Each dot represents a field with an average of 1 to 3 neurons per field. G) The main significant shift in GTP levels is observed in middle age between nTg and 3xTg-AD (p=0.0002). H) No sex differences for free/bound GTP in nTg neurons or I) in 3xTg-AD neurons. N=cultures, each averaged from 6-11 fields with 1-3 neurons per field

### TMRM assay of mitochondrial membrane potential

Cells were incubated with 20 nM TMRM in 0.01 % DMSO (Thermo Fisher, T668; from 100 µM stock in DMSO) in ERB buffer for 30 min at 37°C in ambient CO_2_. TMRM signal was excited with a DPSS laser (561-10 nm, laser power at 0.1) and detected at 566-685 nm (red channel). Colocalization with automatic thresholding (ImageJ/Fiji, v1.53t, NIH) is based on a pixel-by-pixel measure of the Pearson correlation of intensities after automatic subtraction of pixels with zero correlation. Each flame graph contains the pixels from 1-3 neurons of one field. The mean and S.E. are presented for the correlations from ten fields.

### Immunofluorescent staining

After GTP measurements, the neurons attached to glass coverslips were rinsed twice with ice-cold PBS and fixed with ice-cold 4% paraformaldehyde in PBS for 10 min. The fix was removed by one rinse with PBS. Non-specific binding sites were blocked with 2% BSA and cells permeabilized in 0.5 % Triton X-100 in PBS for 1 h at room temperature. Primary antibodies were prepared in 1% BSA, 0.05 % Triton X-100 in PBS and allowed to bind overnight at 4°C, including: anti-Arl8b (1:500; Proteintech, #13049-1-AP), anti-Rab7 (1:350; Millipore, # R8779), anti-Nrf2 (1:100; Santa Cruz Biotechnology, # SC722), anti-NQO1 (1;250; Invitrogen, # 39-3700), anti-mOC78 (1:40; residues 8-11 (SGY), 20-24 (FEV), gift from Charles Glabe [22], and anti-nitrotyrosine (1:100; Millipore, # 05-233). Unbound antibodies were removed by rinsing three times with PBS. Secondary antibodies of goat anti-rabbit Alexaflour 555 (1:1000; Thermo-Fisher, # A31572) and goat anti-mouse Alexaflour 488 (1:1000: Thermo-Fisher, # A11029) were incubated for 1 h at room temperature protected from light. Slips were washed three times with PBS with 5 min between each wash. Nuclei were stained with 1 ng/ml Bisbenzamide (from 1 μg/ml stock; Sigma-Aldrich # B2261) for 2 min in PBS. The stained coverslips were mounted on glass slides using one drop of aqueous mounting medium (PermaFluor # TA-030, Fisher Scientific, CA).

### Statistical analysis

Each experiment was performed at least three independent times (as specified in each figure legend). Results are presented as mean of multiple experiments or all the fields from all experiments and standard error. Statistical significance was determined by using a two-way ANOVA with rejection of the null hypothesis at p<0.05. Follow-up Dunnett’s multiple comparisons and Student’s test was used to determine subgroup significance using Prism software.

## Results

### Age and AD-genotype-related decreases in levels of GTP in hippocampal neurons

GTP is one of the bioenergetic currencies that drives cellular processes essential for vesicle trafficking and clearance, such as endocytosis and autophagy. However, little is known about its fluctuation across age-span and in Alzheimer’s disease mouse model neurons. To explore this, GTP levels were evaluated in primary neurons using the GEVAL biosensor created by Bianchi et al. [20] (Fig.1A). The GEVAL biosensor detects free GTP when excited at 488 nm and bound GTP at 405 nm using a Null probe as a transfection control [21]. GTP is constantly mobilized from the TCA cycle and directly from cytoplasmic ATP to meet free GTP energetic demands of various cellular functions. To test the hypothesis that GTP levels are impaired with age and AD-genotype, we measured the free/bound GTP ratio and bound/free GTP in cultured hippocampal neurons at 14 days in vitro of different ages.

Qualitatively, compared to nTg neurons, a prominent reduction in free-GTP levels occurred in middle age 3xTg-AD (Fig. 1B). Notice the strong overlap of free and bound structures. Bright field images show structurally healthy neurons after transfection to rule out overt cellular damage. By computing the pixel-by-pixel ratio of free GTP/bound GTP (Fig. 1C, D), we mitigate against differences in transfection efficiency. In the non-Tg young control neurons (Fig. 1C), levels of somal and dendritic localized free/bound GTP increased in middle-age before declining sharply in old age neurons. The complementary GTP bound/free images localized to the cell periphery where endocytosis occurs, and accumulate there in old age neurons. For 3xTg-AD neurons in Fig. 1D, young free and bound GTP showed somal and dendritic localization similar to nTg neurons. However, in middle-age a considerable somal shift and decrease in free GTP occurred, which remained low into old-age. In old-age 3xTg-AD neurons, bound-GTP remained localized at the periphery. Therefore, in subsequent quantitative analysis, we focused on free-GTP levels.

Quantitatively, in nTg neurons (Fig. 1E), the levels of free-GTP increased in middle age, followed by a marked decline toward old age (ANOVA F(2, 35)=27.8, p<0.0001). In 3xTg-AD neurons (Fig. 1F), a considerable decrease in free GTP levels occurred at middle-age and remained low in old-age (ANOVA F(2, 46)=11, p<0.0001). Fig. 1G shows comparisons by age and genotype. A marked shift in GTP levels occurs in middle age between both genotypes (ANOVA F(2, 83)=9.6, p=0.0002). The loss of free GTP in middle-age 3xTg-AD neurons is large, while both genotypes exhibit a substantial decrease in free-GTP levels in old-age, suggesting that the cell’s ability to regenerate GTP is restricted by age and genotype. Further subdivision by sex indicates no significant differences for nTg neurons (Fig. 1H) or 3xTg-AD neurons (Fig. 1I). Therefore, variations in GPT levels are independent of the animal’s sex. These results suggest for the first time that GTP levels are modified with aging, showing a large decline in 3xTg-AD neurons in middle-age compared to the rise in nTg neurons before declines in both genotypes in old-age.

### Free-GTP that co-localized in the mitochondria declined in old age

Since GTP is primarily produced in mitochondria by the catalytic function of succinyl-CoA synthetase in the TCA cycle, we explored the presence of free-GTP pools in the mitochondria in hippocampal neurons, by probing for co-localization with the membrane potential dye TMRM, whose positive charge favors distribution into negatively polarized mitochondria [23]. Fig. 2 shows evidence for a strong co-localization between free-GTP and TMRM in young age (Fig. 2Aa-d) and middle age (Fig. 2Ae-h) nTg neurons. In old-age nTg neurons, the colocalization decreased, accompanied by an increase in free-GTP labeled vesicular-shaped structures (Fig. 2Ai-l). In 3xTg-AD, a marked reduction in free-GTP in the mitochondria occurred in middle age (Fig. 2Be-h) and remained low in old age neurons (Fig. 2Bi-l). With age, punctate vesicles were observed outside mitochondria, most prevalent in 3xTg-AD neurons from middle age and old age mice. Fig. 2C shows comparison by age and genotype in the colocalization between free GTP with TMRM (ANOVA genotype F(1,132)=76, p<0.0001; age F(2,132)=47, p<0.0001). In nTg, colocalization increased in middle age (p=0.01) and decreased in old age (p<0.0001). In 3xTg-AD, the correlation decreased in middle and old age (p<0.0001 and 0.0002, respectively). We next determined whether the decline in free GTP could be restored by small molecules targeting bioenergetics and oxidative redox state, followed by vesicular involvement in autophagy.

**Fig. 2.**
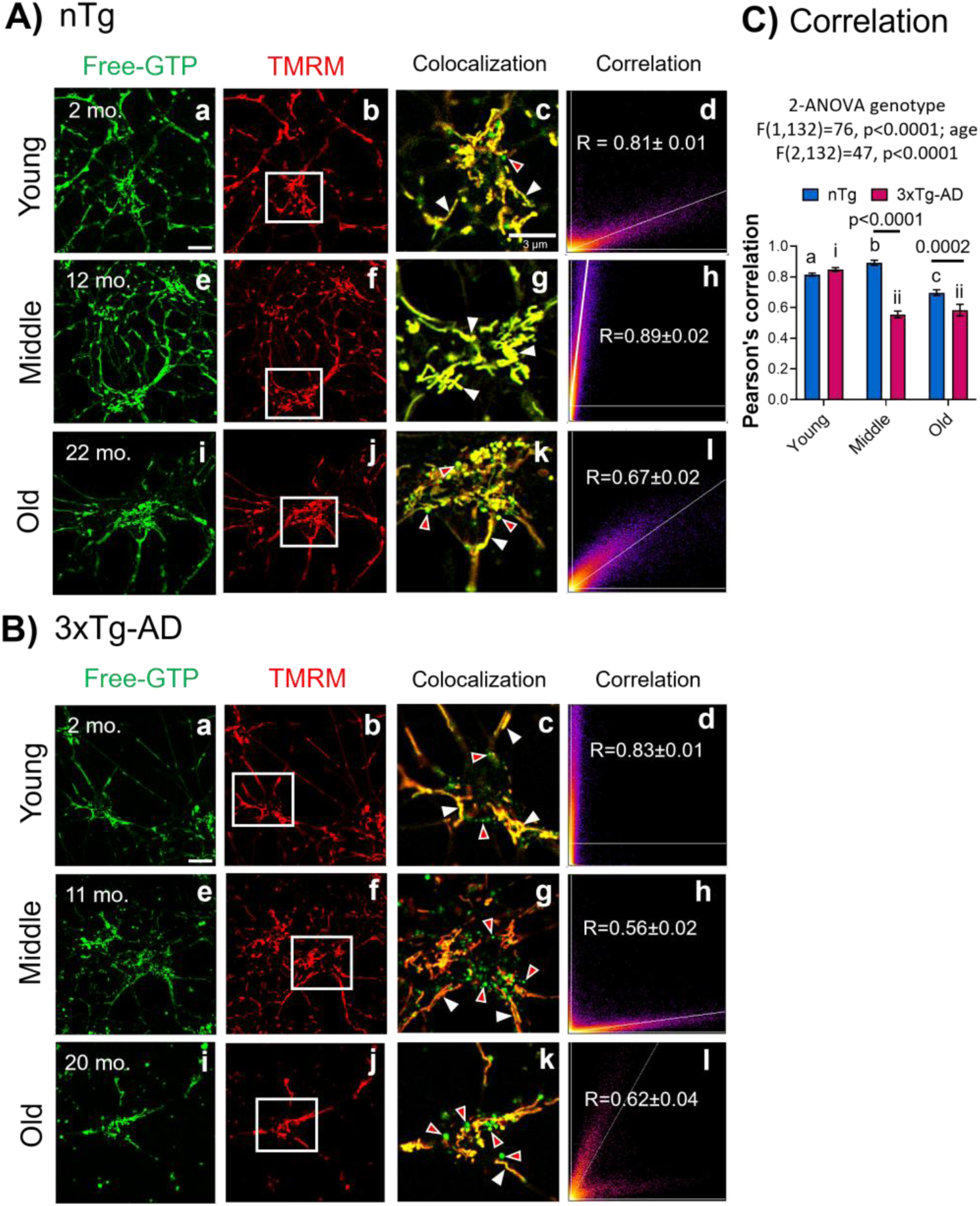
Free-GTP co-localized to mitochondria declined in old age. Hippocampal neurons preloaded with GEVAL free-GTP (488 nm) were incubated with the mitochondrial marker TMRM. Images were merged for yellow pixel level colocalization and quantified (right column) from 10 fields. A) nTg and B) 3xTg-AD neurons by age and the correlation average. Representative images from young neurons (a, b) were merged and magnified in c. Similar representative images for middle-age (e, f) and neurons from old mice (i, j) were merged and magnified in g and k. White arrows indicate co-localized objects between free-GTP and mitochondria. Red arrows indicate green vesicle-shaped structures with free-GTP segregated from mitochondria. Flame maps indicate pixel-by-pixel intensity colocalizations and Pearson’s correlation of co-localization (d,h,l). C) Pearson’s correlation. Different lowercase letters and numbers indicate significant differences after Tukey correction for multiple comparisons. In nTg, free GTP increased the co-localization with TMRM in middle age (b, p=0.01) and decreased in old age (c, p<0.0001). In 3xTg-AD, the correlation decreased in middle and old age (ii, p<0.0001 and 0.0002, respectively). Correlations are mean and S.E. for N=25-30 fields from 3 experiments

### NAD-precursor plus Nrf2 inducer treatment restored GTP levels and size of vesicles in old neurons of both genotypes

Aging, as a major risk factor of AD, is associated with an oxidative redox shift [12, 24, 25] decreases in antioxidant protection [26], and mitochondrial dysfunction [27, 28]. Previously, a lower capacity for maintaining mitochondrial free-NADH was found in old-age neurons from nTg and 3xTg-AD mice [29]. Since the redox energy supply of NAD^+^ is vital to glycolysis and the TCA cycle, forming NADH in the mitochondria, as well as a precursor in the de novo biosynthesis of guanosine monophosphate (GMP), we evaluated nicotinamide as a precursor of NAD^+^ [30] to boost the production of GTP. However, boosting the electron transport chain could also increase the production of oxyradicals. Therefore, we evaluated the combination of nicotinamide with a Nrf2 redox gene inducer, the green tea polyphenol EGCG to control ROS. Rather than finding a single best antioxidant target to support the regulation of age-related decline in redox state, we explore an activator of the cell’s balanced antioxidant/oxidative redox response, the transcription factor Nrf2 (nuclear erythroid factor-like 2). Nrf2 binds at genomic ARE sites (“Anti-oxidant” response elements) to control redox genes [31]. In this study, we use EGCG ((–)-epigallocatechin-3-gallate), the major catechin from green tea [32], as a safe and well-studied Nrf2 activator in combination with nicotinamide to restore the levels of GTP.

In nTg neurons (Fig. 3A), treatment with the combination of nicotinamide and EGCG elevated free GTP in old-age. In 3xTg-AD neurons (Fig. 3B), combined treatment restored free GTP levels in middle-age and old neurons. Fig. 3C, D shows quantitative analysis and a breakdown into treatment with individual drugs. In nTg neurons (Fig. 3C), EGCG alone did not change the free GTP levels, while nicotinamide alone or the combination with EGCG reverted the old age decrements in free GTP to levels of young neurons (2-way ANOVA for age F(2,103)=16, p<0.0001 for treatment F(3,103)=1.6, p=0.2 (subtests in old untreated vs nicotinamide, p<0.0004, and combination p<0.0001). In 3xTg-AD neurons (Fig. 3D), either EGCG alone or the combination of nicotinamide and EGCG reverted middle-age and old neurons to their youthful levels (2-way ANOVA for age F(2,163)=12, p<0.0001; for treatment F(3,163)=7, p=0.001) with indicated subtests.

**Fig. 3.**
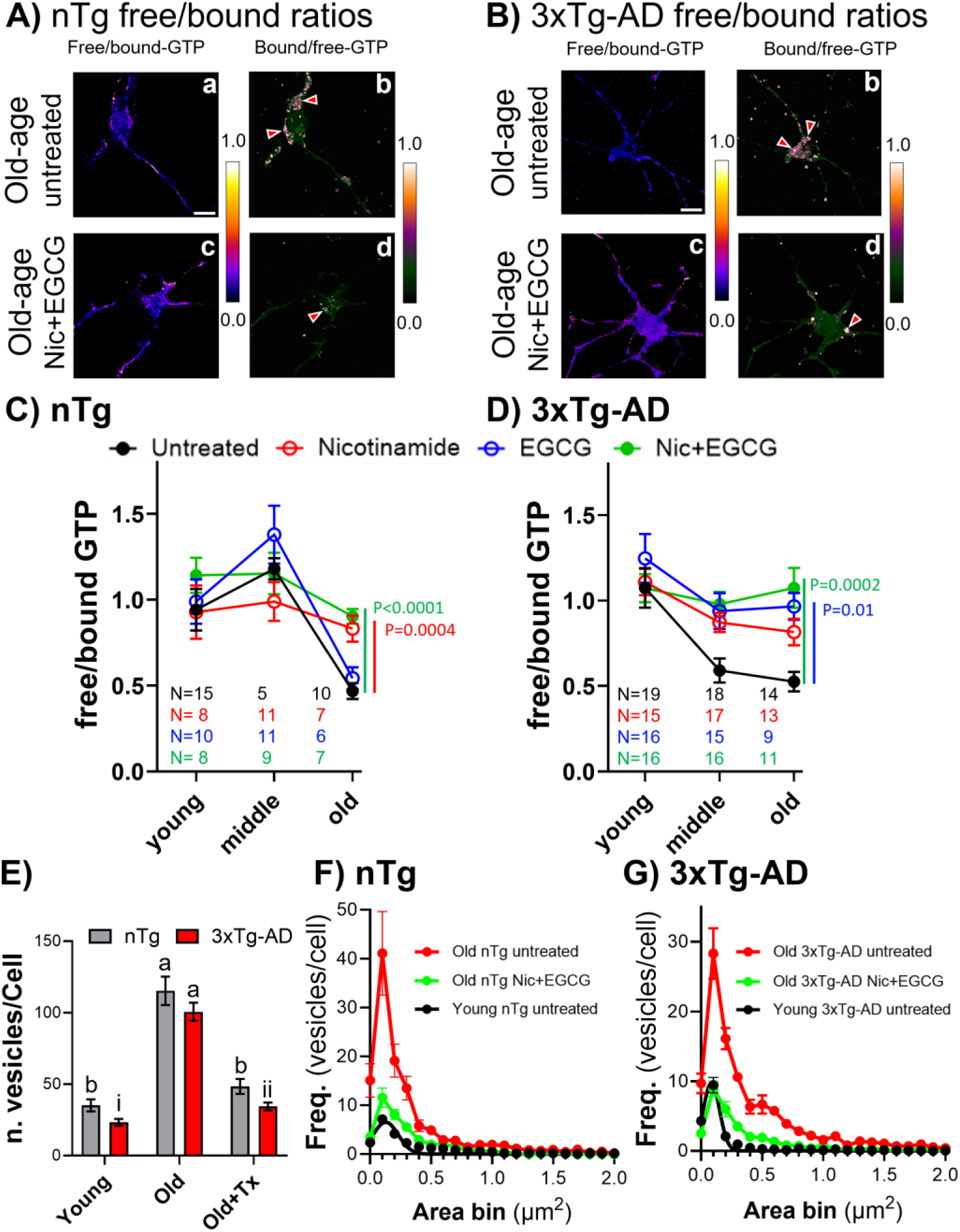
Treatments of old neurons with energy precursor and Nrf2-inducer restored GTP levels and size of vesicles in both genotypes. A) Free/bound GTP levels (a) and bound/free GTP levels (b) in untreated old-age (20 months) nTg neurons. Treatment of old age (23 months) nTg neurons increased the free/bound-GTP levels in the cytoplasm and dendrites (c) and reduced the bound/free GTP levels (d). B) In untreated old age (26 months) 3xTg-AD neurons, Free/bound GTP levels (a) and bound/free GTP levels (b). In old age 3xTg-AD, the decay observed in free/bound GTP levels was restored to basal levels with combination treatment (c), reducing the bound/free GTP levels (d). Red arrows indicate accumulation of vesicles in both genotypes. C) Treatment restored the free/bound-GTP levels in old age nTg neurons. D) In 3xTg-AD neurons, combination treatment prevented the decrease in free/bound-GTP at middle and old ages. Each point is the mean of 7-10 fields from each mouse culture (N cultures). E) A large increase in vesicles with bound GTP was observed in old age in both genotypes. Different lowercase letters and numbers indicate significant differences (p<0.0001) after Tukey correction for multiple comparisons. F and G) Treatment with nicotinamide and EGCG combination reduced the number of bound-GTP vesicles in old-age neurons in both genotypes (nTg: young 2 mo. and old 24 mo.; 3xTg-AD: 2 mo. and 18 mo.)

### Effect of treatment on the number and size of bound-GTP vesicles and rapid Nrf2 activation in 3xTg-AD hippocampal neurons

We hypothesized that impaired vesicular endocytosis was caused by an energy shortage that resulted in a buildup of vesicles and possibly enlargement from failure to complete processing. To explore the mobilization of vesicles corresponding to bound-GTP that accumulated in old age (Fig. 3Ab, 3Bb), the number and size of vesicles following treatment were quantified. In Fig. 3E, a 4 to 5-fold increase in vesicle numbers was observed in old age in both genotypes compared to young age (p<0.0001), with no difference between genotypes. The combination treatment with nicotinamide and EGCG restored the number of old-age vesicles to young levels in both genotypes (2-way ANOVA for treatment F(2,100)=93, p<0.0001; genotype F(1,100)=14, p=0.01, with indicated subtests) (Fig. 3E). Fig. 3F and G show the size distribution in each genotype. The vesicle frequency in old-age nTg neurons decreased by treatment (2-way ANOVA for age-treatment F(2,606)=109, p<0.0001) as well as the size (F(20,607)=44, p<0.0001). In Fig. 3G for old age 3xTg-AD neurons, the frequency and size of vesicles decreased by treatment (2-way ANOVA for age-treatment F(2,567)=65.5, p<0.0001; size F(20,567)=38, p<0.0001).

Prior work showed the transcription of Nrf2 was about 50% lower in middle and old age 3xTg-AD neurons [16]. We next examined the time-course of transcription factor Nrf2 activation as Nrf2 translocation from the cytoplasm to the nucleus following combination treatment (Fig. 4). In Fig. 4A-4C the compartmentalization shifted from the cytoplasm to the nucleus as early as 30 min. as expected for release of Nrf2 from its inactive cytoplasmic binding to KEAP1 and trafficking to the nucleus [33]. Nrf2 stabilization achieved maximal nuclear localization at 30 min that returned to basal levels at 900 min. (15 hr.) (2-way ANOVA for interaction F(4, 41)=6, p=0.0006). In Fig. 4C, the known Nrf2 transcription of a target gene is shown by maximum immunostaining at 30 min. for NQO1, NADH-quinone oxidoreductase, coincident with nuclear localization of Nrf2 in hippocampal neurons.

**Fig. 4.**
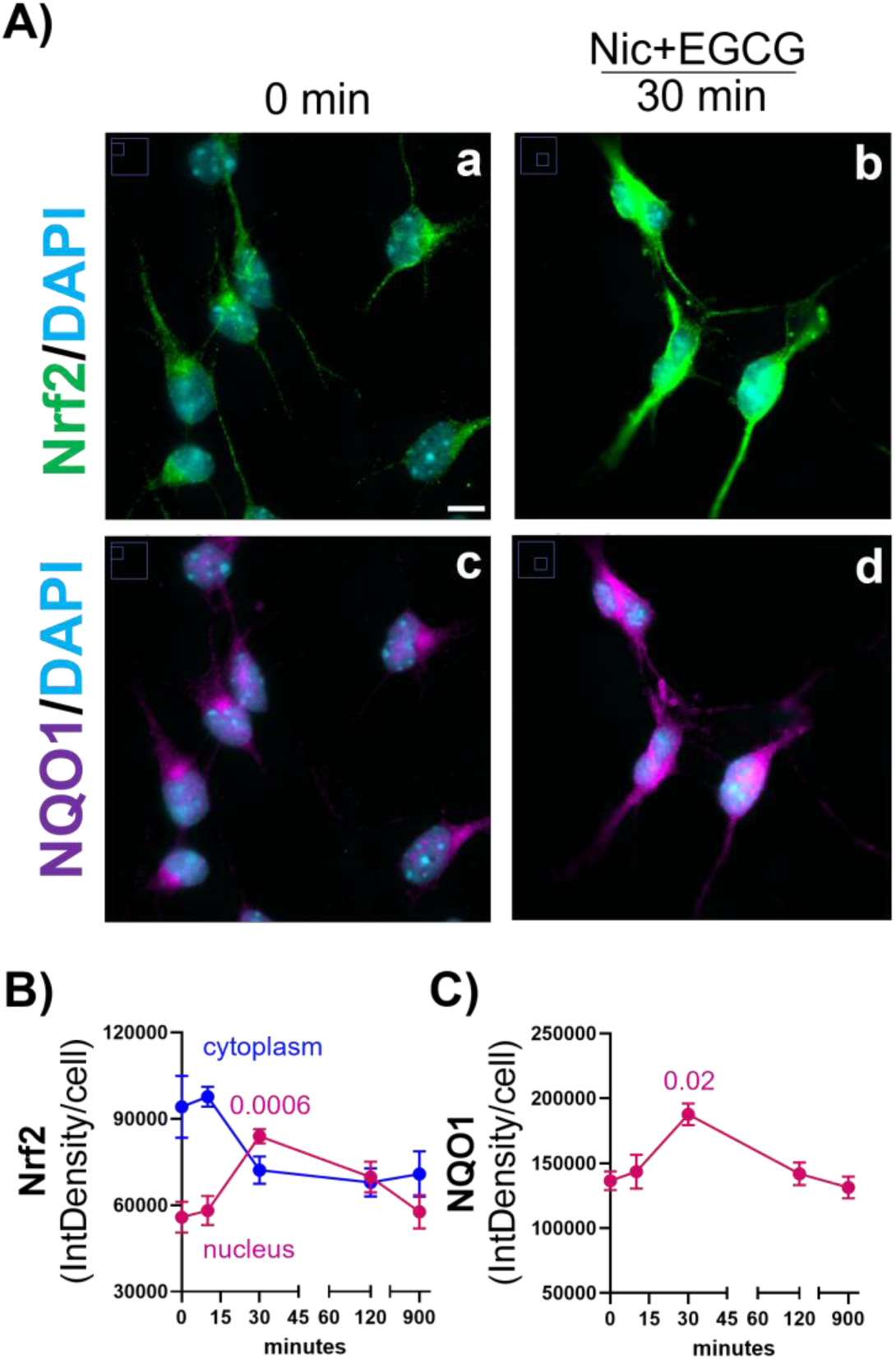
Effect of treatment on rapid Nrf2 activation in 3xTg-AD hippocampal neurons. In A) representative images of cells (a) indicate a rapid Nrf2 immunostained activation at 30 min with treatment in 3xTg-AD hippocampal neurons (b). B) Reciprocal nuclear and cytoplasmic shifts in localized levels of Nrf2 over time. Nuclear Nrf2 levels peaked at 30 min (p=0.0006). C) Temporal induction of NQO1 levels (c) with Nic+EGCG treatment (d) peaked at 30 min (p=0.02)

### GTP is used for autophagy

Our prior work indicated that following endocytosis of the amyloid precursor protein at the cell surface, Aβ accumulated in early and late endosomes as well as autophagic vesicles [1]. Since the processing of endocytic and autophagic vesicles involves many GTPases [7], the age-related GTP deficit might impair autophagic processing. To understand what proportion of GTP consumption is driven by autophagy, we first tested changes in free GTP following inhibition of autophagy using bafilomycin [34]. Fig. 5 shows large increases in free GTP with time of treatment for both nTg (Fig. 5Ad and 5C) (2-way ANOVA for time F(4,26)=4.6, p=0.006; treatment F(2, 26)=55, p<0.0001) with indicated subtests) and 3xTg-AD (Fig. 5Bd and 5D) (2-way ANOVA for time F(4,27)=35, p<0.0001; treatment F(2, 27)=100, p<0.0001) with indicated subtests). To determine if the neurons had enough capacity to maintain free GTP levels while autophagy was stimulated, we treated neurons with rapamycin to induce autophagy [35]. Fig. 5Af and 5C show that autophagy induction caused a net consumption of GTP in nTg neurons. In 3xTg-AD neurons, rapamycin failed to affect free GTP levels, either because it was unable to stimulate autophagy or that the cells were able to handle the extra load and maintained GTP levels (Fig. 5Bf and 5D). Together, these results suggest that GTP is heavily used in autophagy and that 3xTg-AD neurons lose the capacity to respond to autophagic induction.

**Fig. 5.**
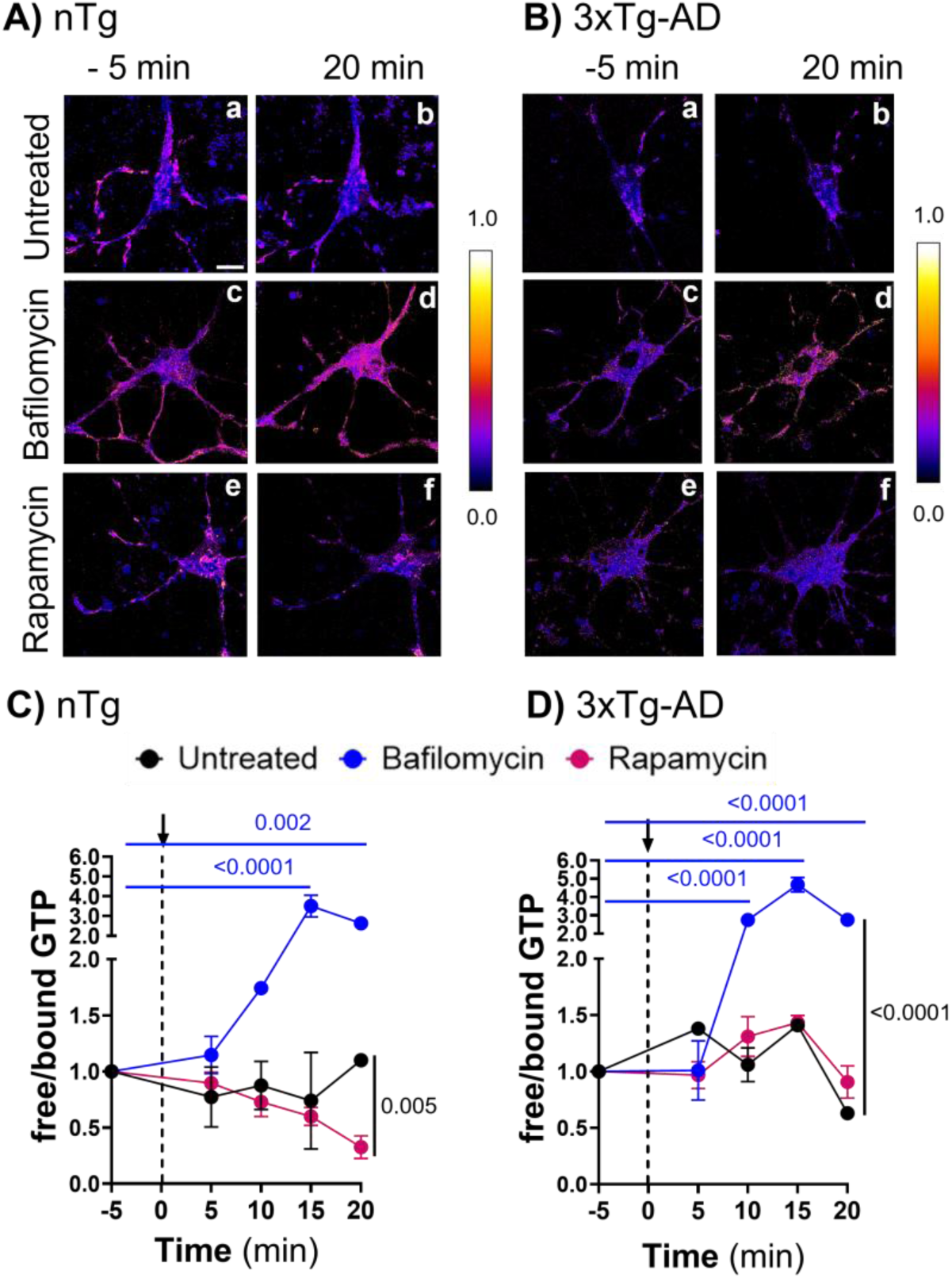
GTP used for autophagy. A and B) Representative images of the ratio of free/bound from nTg (Aa, Ac, and Ae) and 3xTg (Ba, Bc, and Be) neurons from middle age (12 mo. male) before treatments at –5 min (base line), and 20 min after treatment: untreated (Ab, Bb), 75 nM Bafilomycin (Ad, Bd) and 20 nM Rapamycin (Af, Bf). Free GTP accumulated in nTg middle-age neurons treated with bafilomycin (D and Ad) and 3xTg middle-age neurons (D and Bd). Net GTP was consumed by induction of autophagy with rapamycin only in nTg neurons (C and Af). The arrow indicates the time of treatment addition. Some error bars are smaller than the symbol. N=3-5 fields with 1-2 cells/field/point and condition (Representative experiment of 3)

### Accumulation with age of vesicular GTPases involved in endocytosis (Rab7) and lysosomal degradation (Arl8b)

Small GTPases drive essential processes to clear aggregated or misfolded proteins in the cell by the process of autophagy [7]. Since proper functioning of GTPases requires an adequate supply of GTP, impaired GTP-dependent vesicular trafficking is likely to accumulate vesicles at incomplete stages of processing. To explore impairments in autophagy with age, after measuring GTP levels in neurons from mice across the age-span, cells were fixed and immunostained for GTPases Rab7, a marker of late-endosomes [36], and Arl8b, a lysosomal protein [37]. Fig. 6A-4H shows a marked accumulation of Rab7 and Arl8 in the axons and dendrites of old age nTg neurons (Fig. 6Ac) that was enhanced in 3xTg-AD neurons (Fig. 6Cc) for Rab7 (2-way ANOVA for time age F(2,330)=36, p<0.0001). Supporting these results, earlier findings of an overlap between Rab7 and intracellular Aβ (iAβ) indicated an accumulation of iAβ-loaded late-endosome-marked vesicles in 3xTg-AD neurons [1].

**Fig. 6.**
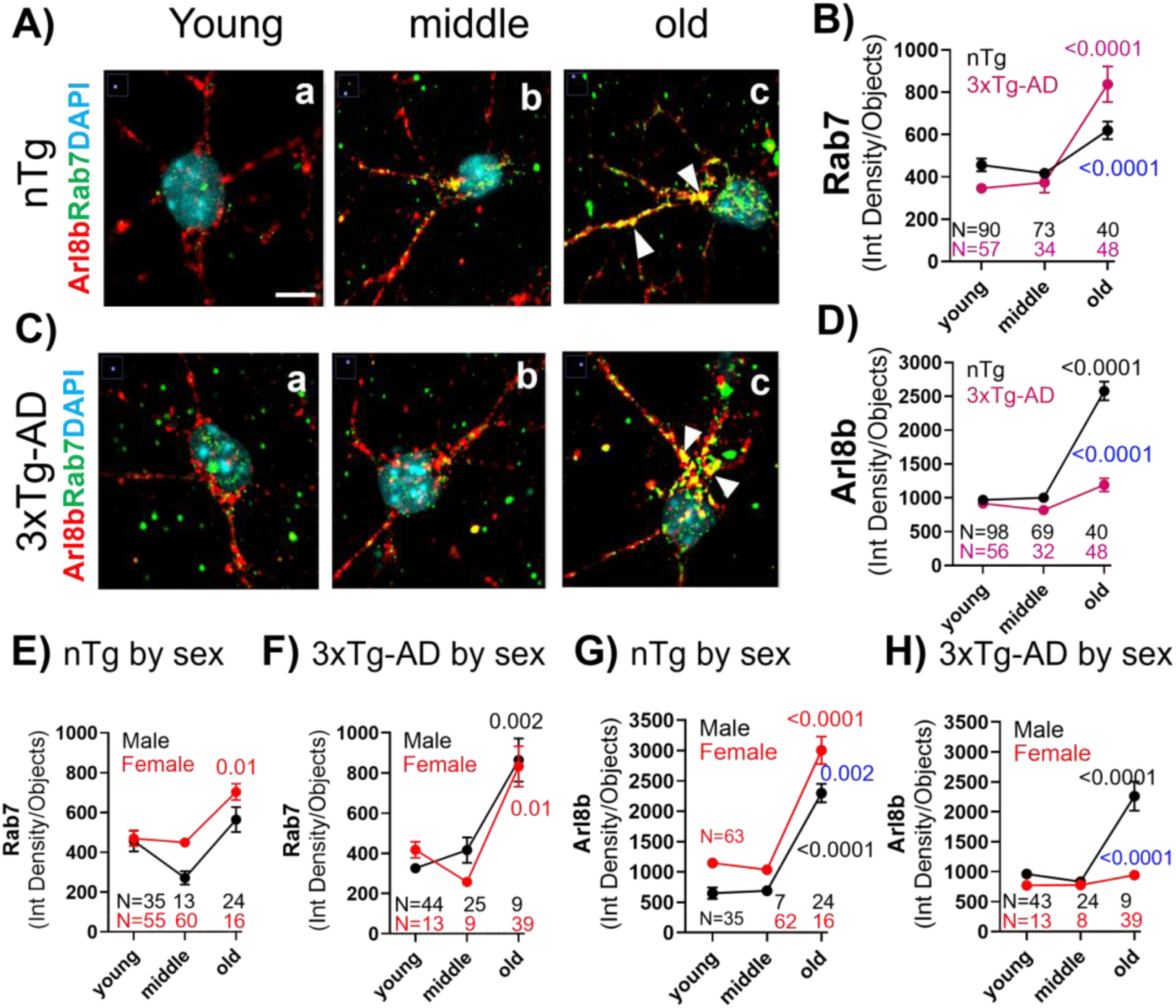
Accumulation with age of vesicular GTPases involved in endocytosis (Rab7) and lysosomal degradation (Arl8b). A and C) Representative images of cells staining for Rab7, Arl8b, and DAPI in nTg (Aa-Ac) and 3xTg-AD (Ca-Cc) at different ages. White arrows indicate prominent colocalized accumulation of Rab7 and Arl8b in old age in axons and dendrites (Ac and BC). B) Quantitative analysis of greater Rab7 accumulation in old 3xTg-AD than old nTg neurons. Each dot is the mean of fields with 10-25 cells in N fields from multiple cultures. D) Quantitative analysis of greater Arl8b accumulation. Larger accumulation of Arl8b in old nTg neurons than 3xTg-AD. E) Female nTg neurons accumulated more Rab7 vesicles than male in middle and old age neurons. G) Female nTg neurons accumulate more Arl8b than male neurons. F) No sex differences in Rab7 accumulation in the old 3xTg-AD neurons. H) Larger old male 3xTg-AD accumulation of Arl8b than female. Blue P values in D, F, I and J indicate differences in old age

These results suggest a fusion between the late-endosome (Rab7) and lysosome (Arl8b) in the neurites of hippocampal neurons, and deficient endocytic degradation. Statistical analysis in Fig. 6B shows a marked increase in Rab7 levels in old age in 3xTg-AD neurons, while a smaller increase was observed in old age nTg (2-way ANOVA for time age (F(2,330)=36, p<0.0001). Fig. 6E shows a greater Rab7 accumulation in female old age nTg mice than males (2-way ANOVA for age F(2,191)=13, p<0.0001; sex F(1, 191)=7, p=0.009). In old age 3xTg-AD neurons, Fig. 6F shows a similar tendency in Rab7 accumulation in both sexes (2-way ANOVA for age F(2,131)=16, p<0.0001; sex F(1, 131)=0.2, p=0.67). In Fig. 6D, Arl8b accumulated in old nTg neurons but failed to increase in old 3xTg-AD neurons (2-way ANOVA for age F(2,337)=89, p<0.0001; genotype F(1, 337)=68, p<0.0001). A greater sex-dependent increase in Arl8b was observed in nTg neurons compared to male neurons (Fig. 6G). The failure to increase Arl8b levels in old 3xTg-AD neurons, was primarily among female neurons (2-way ANOVA for age F(2,130)=38, p<0.0001); sex F(1,130)=39, p<0.0001) (Fig. 6H). These results suggest that aged cells have alterations in the vesicular clearance mechanisms driven by Rab7 and Arl8b GTPases, which could be caused by the bioenergetic deficits of intracellular free-GTP in old age.

### Bioenergetic boost with nicotinamide and redox protection with EGCG restored Rab7 and Arl8b to lower levels in both genotypes

Once we confirmed that age-related GTP deficits were mitigated with nicotinamide and EGCG treatment, we explored whether the accumulation of late endosome vesicles labeled with antibodies to the Rab7 GTPase or lysosomal vesicles labeled with antibodies to the Arl8b GTPase were reduced in response to the proposed bioenergetic-redox boost. Either the accumulation of GTPase-labeled vesicles is a protective response for which bioenergetic treatment could boost protection or the buildup is pathologic and could be cleared with increased energy supplies. Fig. 7A, B suggest that levels of Rab7 and Arl8b at young ages in either genotype were unchanged by treatment with nicotinamide and EGCG. However, treatment of old neurons of either genotype with nicotinamide and EGCG lowered levels of Rab7– and Arl8b-stained vesicles. Indications of colocalization suggest initiation of a fusion process that is unable to be completed in old age neurons. Statistically, in Fig. 7C, each drug or the combination treatment lowered Rab7 vesicle staining in old neurons to young nTg levels (2-way ANOVA for age F(2,693)=3, p=0.05; treatment F(1, 693)=4.3, p=0.004). In 3xTg-AD neurons, Fig. 7D shows efficacy of each drug in lowering the old-age rise in Rab7 vesicles, but the combination was most effective in lowering late endosome levels in old neurons to young levels (2-way ANOVA for age F(2,489)=22, p<0.0001; treatment F(3, 330)=5, p=0.001). For Arl8b-stained vesicles in old age nTg neurons (Fig. 7E), either nicotinamide or the combination with EGCG partially reversed the elevated levels of vesicles (2-way ANOVA for age F(2,700)153, p<0.0001; treatment F(1, 700)=6, p=0.0004). In old 3xTg-AD neurons (Fig. 7F), either drug alone or the combination lowered the old-age Arl8b levels to those of young neurons (2-way ANOVA for age F(2,482)=12, p<0.0001; treatment F(3, 482)=9.8, p<0.0001). These treatment results suggest that the old age increases in Rab7-labeled late endosome and Arl8b-labeled lysosomes are not protective but suffer from insufficient energy to complete the processes of vesicular autophagy via lysosomal degradation. We next examined whether this affected the intracellular accumulation of Aβ.

**Fig. 7.**
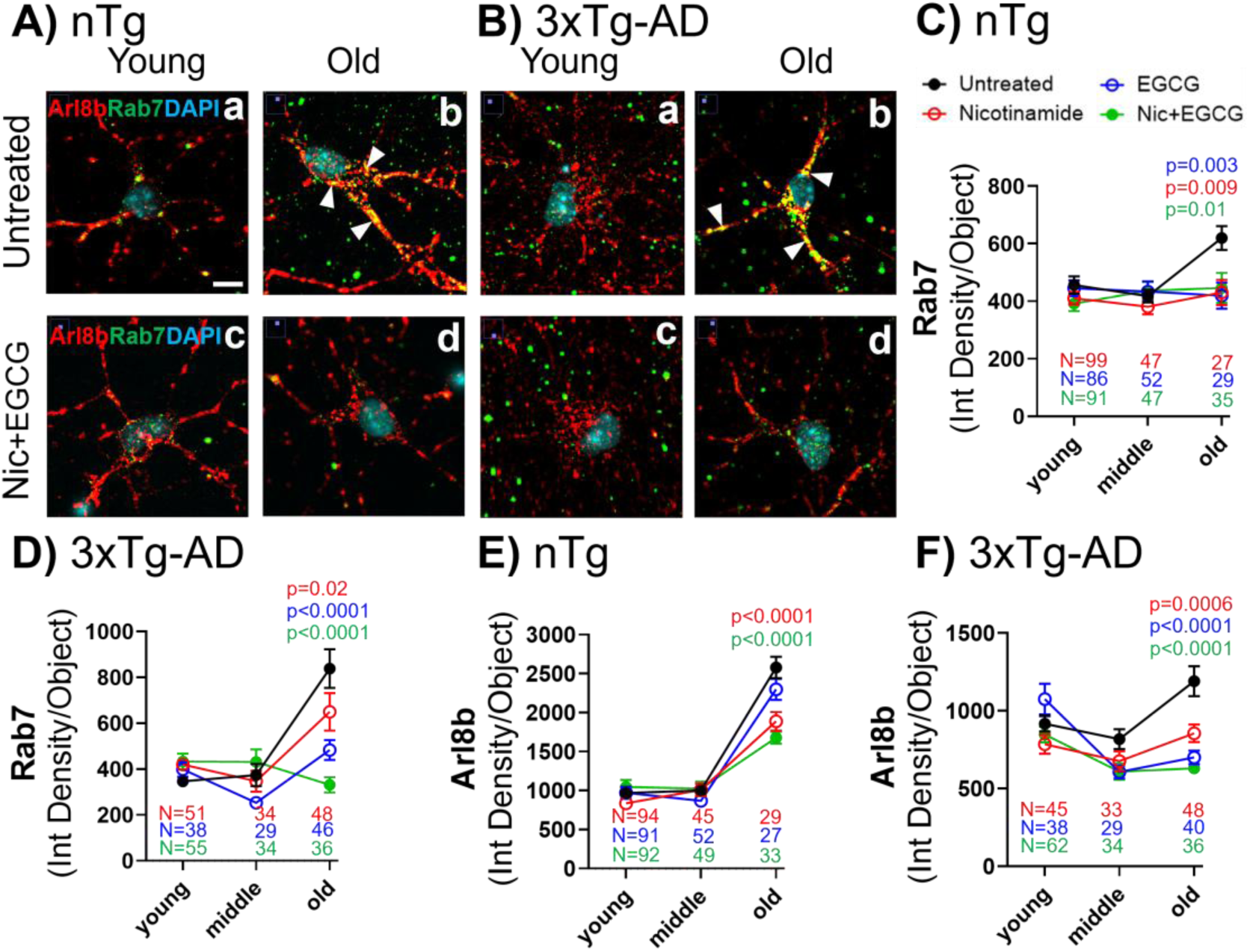
Bioenergetic boost with nicotinamide and redox protection with EGCG lowered Rab7 and Arl8b levels in old-age in both genotypes. A) In nTg neurons, and B) 3xTg-AD neurons, co-localization between Rab7/Arl8b increased in axons and dendrites in old-age (b). Treatment with nicotinamide (Nic) + EGCG reduced both Rab7 and Arl8b signals and their co-localization only in old age neurons (d). White-arrows indicate the co-colocalization in the cell. No co-localization signal was observed in young-age in either genotype. C) In nTg old neurons, the Rab7 increase was remediated by nicotinamide, EGCG or the combination (colors indicate p values). D) In old-age 3xTg-AD neurons, the Rab7 increase in old neurons was partially reversed by nicotinamide, more so by EGCG or the combination. E) The Arl8b increase in nTg old neurons was reduced by nicotinamide, or the combination with EGCG. F) In old-age 3xTg-AD neurons, the Arl8b increase in old neurons was reversed by nicotinamide, EGCG or the combination. Each dot is the mean of n fields containing 1-5 cells

### Nicotinamide and EGCG reduce Aβ accumulation and protein oxidation in aged 3xTg-AD hippocampal neurons

Since intracellular Aβ aggregates increase across the lifespan of the 3xTg-AD model in vivo (Pontrello [38] 2022), and in vitro [1] and a hallmark for brain aging and the progression of AD is protein oxidation, including tyrosine nitration [1,39], we immunostained neurons for Aβ accumulation and protein nitration and evaluated the effects of treatment with nicotinamide and EGCG. Compared to nTg neurons, Fig. 8A shows large intracellular Aβ aggregates in old-age 3xTg-AD neurons detected by binding of an Aβ aggregation-specific antibody mOC78 (Fig. 8Ac) [38, 22]. As expected, Aβ aggregates in old-nTg neurons were negligible (Fig. 8Aa). These results replicate the intracellular Aβ deposits previously reported that began in middle-age 3xTg-AD in vitro [1], and in vivo [38]. In Fig. 8Ca and 8Cc age-related deposits of nitrotyrosine tagged aggregated proteins in the soma and dendrites in both genotypes. The localization of these oxidized protein deposits appears non-coincident to those of Aβ aggregation, suggesting a separate age-related pathway for oxidation of protein. Combination treatment with nicotinamide and EGCG for just 16 hr. (Fig. 8Ad and 8B) reduced Aβ-aggregate deposition in 3xTg-AD hippocampal neurons close to levels in nTg neurons (2-way ANOVA genotype F(1,40)=6.6, p=0.01; treatment F(3, 40)=4.4, p=0.009). The combination of nicotinamide and EGCG tended to be more effective than either alone (p=0.0001). Similarly, Fig. 8Cd and 8D show that protein nitration in old-age neurons was reduced by nicotinamide or EGCG or the combination in 3xTg-AD neurons (ANOVA treatment F(3,26)=5.7, p=0.003), but only nicotinamide and combination reduced the protein nitration in old age nTg neurons (ANOVA treatment F(3,15)=3.8, p=0.03). Together, these results suggest that treatment provides the energy and redox balance needed to complete Aβ processing by endocytosis, lysosomal fusion and autophagy.

**Fig. 8.**
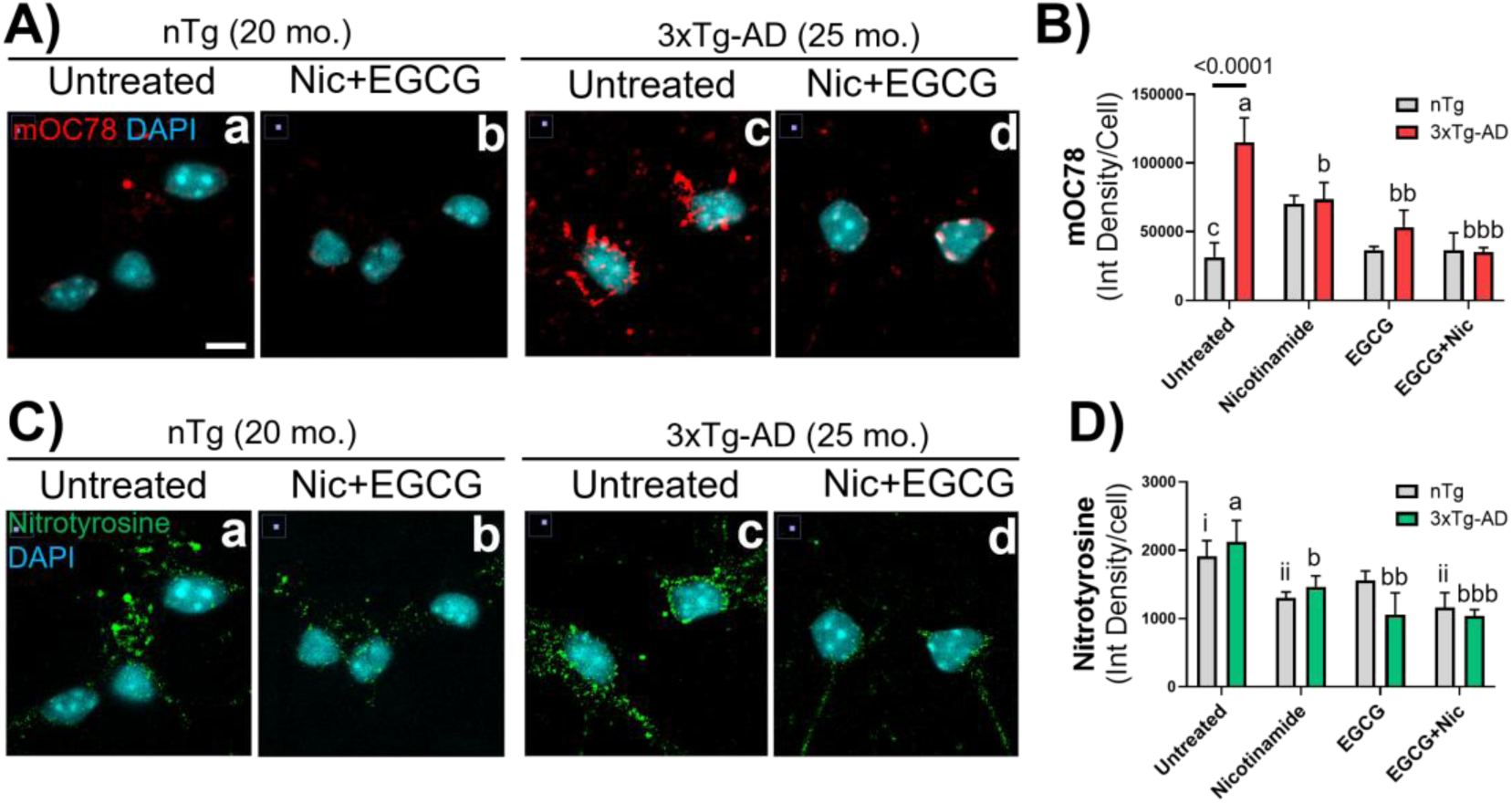
Nicotinamide and EGCG reduced Aβ accumulation and protein oxidation in aged 3xTg-AD hippocampal neurons compared to non-transgenic neurons. A) Aβ aggregates in the cytoplasm of old-3xTg-AD neurons (25 mo.) were cleared following combination treatment. B) Quantitatively treatment with nicotinamide (b, p=0.04), EGCG (bb, p=0.01) or the combination (bbb, p=0.0001) reduced the Aβ accumulation. Treatment of old-nTg neurons (20 mo.) caused no changes in mOC78 levels.C) An extensive protein nitrosive oxidation occurs in old age in both genotypes (Ca,c), that is largely reversed by combination treatment (Cb, d). D) Quantitative analysis shows reduced levels of nitrotyrosine following treatment with nicotinamide, EGCG or the combination. D) Tyrosine nitration was reduced by treatment with nicotinamide (b, p=0.03), EGCG (bb, p=0.008), or combination (bbb, p=0.002) in 3xTg-AD neurons, as well as with nicotinamide (ii, p=0.04) and combination (iii, p=0.03) in old-nTg neurons. Different letters indicate significant differences after Tukey and Dunnett correction for multiple comparisons

## Discussion

### Autophagic impairment with age and AD

Autophagy plays a crucial role in cellular proteostasis [40]. However, what about aging and bioenergetics cause impairment in the detailed molecular mechanisms leading to autophagic failure are poorly understood. In this study, we proposed that aging limits the energetic capacity needed to power autophagy and that the bioenergetic capacity of the AD genotype is further compromised. We report an age-dependent reduction in ratiometric measurements of free/bound GTP levels in living hippocampal neurons, which takes place mainly in mitochondria. Our results demonstrate that the reduction in free-GTP levels occurs in old age in nTg neurons and is accelerated in middle age for 3xTg-AD neurons. This earlier loss of free GPT colocalized with mitochondria in 3xTg-AD neurons could be a key component in autophagic impairments in AD.

Emerging evidence shows that suppression of mitophagy drives cellular aging phenotypes in a PINK1/Parkin/p62-dependent pathway [41]. Following senescence induction by radiation, mitophagy becomes impaired. Loss of mitochondrial turnover driven by autophagy contributes to an accumulation of dysfunctional mitochondria, and an inflammatory DAMP response [42] as a crucial feature of senescent cells. This scenario could result from decrements in mitochondrial bioenergetics, including a decreased supply of GTP for autophagic vesicular processing.

We show that free-GTP levels are sensitive to pharmacological induction and inhibition of autophagy in nTg hippocampal neurons. Induction with rapamycin increased free-GTP consumption indicating reserve autophagic capacity that was more limited in 3xTg-AD neurons. This suggests that the AD neurons were already operating at maximum capacity because of the amyloid load. It has been reported that hippocampal accumulation of mAPP and Aβ impairs mitochondrial dynamics, biogenesis, synaptic activity, glucose metabolism, and autophagy, leading to neuronal dysfunction [43, 44] . In our study, we also found large intracellular Aβ aggregates localized in the cytoplasm in old age 3xTg-AD hippocampal neurons, as well as in the brains of these mice [38]. Previously, we detected intracellular Aβ aggregates (iAβ) in the pathways responsible for vesicular trafficking, including endocytosis and autophagy in old adult neurons from 3xTg-AD model mice [1]. Accumulation of Aβ aggregates and longer Aβs in Rab7-positive late endosomes and cathepsin D-positive autophagolysosome and mitochondria suggest that age-related changes in iAβ processing limits completion of autophagic and endocytic processes in aged neurons [1].

High colocalization of free-GTP with TMRM at young-age indicate a major production site in mitochondria, but also associated with old age enlarged cytoplasmic vesicles. The mitochondrial GTP is probably made by the catalytic function of succinyl-CoA synthetase in the TCA cycle in the matrix or by nucleoside phosphate kinase (NME4) in the mitochondrial intermembrane space [45]. However, as age progresses, a decrease in GTP colocalization with mitochondria in old age indicates an age-related decline in GTP production by these enzymes and a possible shift to cytoplasmic NME1 and NME2 catalysis of ATP to GTP [46] localized near enlarged vesicles, as previously seen for Rab7 immunostaining [1] or invasive filopodia [47]. The enlargement could be due to insufficient GTP levels needed to complete autophagy. These lower GTP levels could be attributed to decreased mitochondrial production of ATP due to Aβ inhibition [44]. Another possibility is that the stress of pathology already strains the maximal capacity for ATP and GTP generation. Since autophagic vesicular processing is GTP-dependent [7], an age-dependent reduction in the GTP levels may cause inefficient vesicular autophagic clearance.

### Specific small GTPases

We report clearance of Arl8b and Rab7 labeled vesicles in nTg and 3xTg-AD old age following boosting bioenergetics and redox balance. Early endosomal and lysosomal abnormalities in AD brains, precede the appearance of extracellular amyloid plaques, and are related to alterations of γ-secretase function and APP metabolism, producing longer Aβ peptides in model cells [48] and in human brain [49]. In our study, a marked accumulation of the colocalization of bound/free GTP with small GTPases Arl8b and Rab7 in both genotypes suggests a lack of endolysosomal mobilization in old age neurons.

The small GTPase Arl8b localizes in the GTP-bound form to the lysosome and mediates the bidirectional transport of lysosomes on microtubules, playing a critical role in lysosomal activity [50, 51]. Boeddrich et al. [52] reported positive correlations of Arl8b levels with Aβ in the 5xTg-AD mouse and abnormal aggregation of Arl8b in AD brains. Arl8b in the lysosome is required for late endosome-lysosome fusion, dependent on, GTP/GDP cycling. The large Arl8b-labeled vesicles that we see could be lipid droplets that accumulate in AD-like neurons [53, 54]. Mobilization of lysosomal-related vesicles into the axon is dependent on proper coupling of the adapter system that includes the BORC complex, the small GTPase Arl8b, and the adapter protein SKIP, which mediates coupling of lysosomes to kinesin-1 [51]. Although we found no sex differences in GTP depletion or Rab7 levels with age or AD-genotype, Arl8b levels in 3xTg-AD remained low in females at all ages but rose sharply in males. This suggests sexual dimorphism at specific points in the autophagic pathway in old age such that females maintain their lysosome levels of Arl8.

Disturbances in the BORCs system leads to axonal degeneration characterized by an accumulation of autophagosomes and axonal swellings filled with autophagosomes and small mitochondria [55]. De pace et al. [55] describe axonal swellings heavily stained for Tau that exhibit an aberrant swirl-like organization of microtubules. Mobilization of vesicles positive to Arl8b and Rab7 colocalization could influence a reduction of p-Tau accumulation, avoiding congestion of vesicular trafficking. p-Tau clearance contributes to proper vesicular transport along microtubules, favoring the mobilization of lysosomes and autophagolysosomes. Previously, nicotinamide mononucleotide treatment reduced the Aβ-induced accumulation of p-Tau in CA1 of mouse brains [56]. Also of relevance are the mitochondrial alterations and lysosome dysfunction seen in human fibroblasts from sporadic AD patients [57]. Interestingly, treatment of these cells with the NAD precursor nicotinamide riboside lowered ROS levels but did not restore lysosomal autophaghic function.

Aβ processing failures dysregulate endosomal trafficking in mouse models of AD [7]. Extensive Aβ accumulation is observed in early– and late-endosomes in hippocampal neurons [1]. Cycles of the active and inactive GTP-bound states of Rab7 affect the lysosome regulation of mitochondrial networks [58]. We report an age-related accumulative pattern of late endosomes in both genotypes, which could be related to a deficiency in the turnover of the bound GTP state of Rab7 due to low GTP levels. A marked colocalization of Rab7 with Arl8b indicates stalling of endolysosomes filled with Aβ aggregates, hindering completion of the autophagic vesicular trafficking. A functional Rab7 is required for the normal development of the autophagy pathway [59]. In our study, a sex comparison between males and females shows a slight increase in Rab7 immunoreactivity in females nTg [1]. These results support a sex factor in the iAβ accumulation for mOC78 immunoreactivity [38] contributing to the increased risk of AD among women.

GTP deficits with age and the corresponding deficits in substrate availability for optimal small GTPase function leads to failures in autophagy with inefficient clearance of protein aggregates, including Aβ. Our study suggests that decreased GTP levels may be a limiting factor in the functioning of Arl8b, Rab7 and other small GTPases. This age-related incomplete vesicular mobilization could be targeted for pharmacological interventions to promote autophagic and endosomal pathways [49].

### Combined bioenergetic and redox treatment

Aging is a major risk factor in AD, but the cause of aging or the age-related triggers for AD pathology are unknown [60]. Dozens of mechanisms have been proposed for this complex syndrome. The complexity of AD may require a combination of treatments to target more than one mechanism. Aging is associated with an oxidative redox shift, which contributes to overproduction of ROS linked to mitochondrial dysfunction [12, 61]. We hypothesized that the redox-bioenergetic impairment of aging contributes to AD pathogenesis, including the failure of autophagy to complete the removal of Aβ aggregates. Age-related reduction of mitochondrial function is accompanied by a decline in the mitochondrial capacity for ATP production by oxidative phosphorylation [62], a reduction in the NAD^+^ pool [63] by consuming processes (PARP, CD38, and SIRT1) [64], and further an increase in ROS generation by an inefficient electron transport chain (ETC) [26]. NAD^+^ depletion triggers cell death via mitochondrial depolarization and mediates cytotoxicity in human neurons with impaired autophagy [65]. We restored the age-related decline in free-GTP levels in primary hippocampal neurons of old age by treatment with an NAD-precursor nicotinamide in just 16 h. Nicotinamide likely serves as the substrate for NAMPT and MNNAT to regenerate NAD^+^ in the salvage pathway. NAD^+^ is also required for de novo synthesis of GMP (and GTP) via inosine 5‘-monophosphate dehydrogenase (IMPDH) catalysis of the IMP to XMP [66]. We are investigating the participation of NME isoforms in rapid localized synthesis of GTP from ATP [46, 67] because of their involvement in GTP-driven vesicular trafficking along microtubules [68], in aging, and AD. Supporting our results, pharmacological interventions with other NAD^+^ precursors including nicotinamide riboside (NR), nicotinamide mononucleotide (NMN), and stimulation of autophagy with rapamycin show anti-senescence properties and autophagic reactivation [69, 70]. In this context, NMN promotes the expression of proteins related to autophagosome formation, including Beclin-1 and LC3II in AD mice [56]. These results suggest a bioenergetic GTP deficit in aging, exacerbated in our AD model that can be remediated by NAD-precursor supplementation. However, NAD-precursor treatment alone failed to raise the decreased levels of GSH seen in 3xTg-AD neurons, as much as a combination of nicotinamide and a redox-regulating Nrf2-inducer was able to restore GSH above nTg levels [71].

### Balanced redox control by Nrf2 inducers

In this work, we demonstrate a rapid activation of Nrf2 by EGCG plus nicotinamide to induce a balanced redox response, including NQO1. This confirms our previous finding of induction of γ-glutamyl cysteine ligase (GCLC) with another Nrf2 inducer, 18-α-glycyrrhetinic acid [71], contributing to induction of mitochondrial catalase, HO-1, malic enzyme and glutathione handling (GST, thioredoxin reductase, GCLC). Other electrophiles such as sulforaphane activate Nrf2 and mitigate redox stress [72]. Since nicotinamide-boosting could increase the production of oxyradicals from the ETC, we treated the hippocampal neurons with EGCG to control an oxidative redox state that causes ROS overproduction. However, emerging evidence points to low ROS levels as a trigger for induction of autophagy [41]. This information suggests that fine regulation of the redox system is key to cellular proteostasis. Other possible targets of EGCG could be NMNAT2 to promote the synthesis of NAD from nicotinamide [73], tau clearance [74], and inhibition of Aβ oligomerization [75].

### Age-related ROS levels

Oxidation of macromolecules also participates in aging and age– and AD-related decline of mitochondrial function [12, 76]. Notably, a proteomic approach identified a set of nitrated proteins, including α-enolase, H^+^-transporting ATPase, and peroxiredoxin-2 in the brains of both Mild Cognitive Impairment (MCI) and AD patients [77]. In our work, reduction of tyrosine nitration by either nicotinamide or EGCG treatment in old age neurons suggests a combination antioxidant effect. Previously, nicotinamide mononucleotide (NMN) treatment has been associated with its ability to activate the Nrf2/keap1/NQO1 pathway, inhibiting Aβ-induced oxidative damage in AD mice [56]. Regulation of autophagy related to nicotinamide treatment also involves Nrf2 activation-driven modulation of p62 autophagy protein levels [56]. Due to the age-related leakage of ROS from mitochondrial complexes and the TCA cycle [78], coupled with the ROS from NAD(P)H oxidoreductases [2] and Aβ processing [79], we predict that GTP levels may shift reductive from an oxidative environment which was remediated by Nrf2 induction with EGCG. Nrf2 activation also upregulates glucose uptake that could benefit old and AD neurons [80].

### Limitations and future studies

We used an in vitro monolayer of neurons to model metabolism dysfunction, which may not reflect the complexity of extracellular matrices in an in vivo environment. However, that we find large effects in the absence of an aging immune system, blood supply and hormones suggests that some effects are intrinsic to the aging neurons themselves. The reversal of age-related phenotypes in less than 24 hr. by energy-precursor and Nrf2-inducer suggests direct metabolic and epigenetic limitations in age-related neurophysiology. Although, compared to younger ages and nTg neurons, 3xTg-AD neurons cultured in vitro under constant and controlled conditions have a defined genotype with human APP (SWE), PS1 (M146V), and Tau (P301L) transgenes to mimic neuropathological features of familial AD, they are limited in application because of the relative overproduction of APP in this model. All cell models do not provide influence from environmental effects on epigenetic changes to the genotype including dietary changes, sleep, or physical or mental fatigue that occur in AD. Despite these limitations, in vivo dosing of nicotinamide with EGCG could be studied for target engagement in the mouse hippocampus and for cognitive improvement.

### Conclusions

This study provides evidence of age– and AD-related GTP level impairments using a genetically encoded GTP sensor in hippocampal neurons. These impairments drive autophagic and endosomal abnormalities. The additional genetic burden of AD genes in the 3xTg-AD hippocampal neurons accelerated the decline in free-GTP levels in middle age. GTP substrate restrictions for small GTPases related to lysosomal and endosomal trafficking may impair the completion of the autophagic process for clearance of Aβ and other damaged proteins. Treatment for just 16 hr. with nicotinamide as an NAD precursor, in combination with EGCG as a Nrf2 inducer, restored the decline in GTP levels in advanced age in both genotypes, restored autophagic clearance of Aβ and reversed levels of oxidized proteins to young age levels. Boosting energy and redox homeostasis are promising strategies to rescue hippocampal neurons from bioenergetic deficits in aging and earlier in AD.

## Author’s contributions

GJB secured funding. RAS and GJB designed the experiments. RAS and JMM performed the experiments. GJB and RAS wrote the first draft. All authors contributed to editing and approved the final version of the manuscript.

## Acknowledgments

We thank Dr. Mikhail A. Nikiforov for the GTP evaluator (GEVAL) plasmids and the UC Irvine Laboratory for Fluorescence Dynamics for use of their confocal microscope under NIH P41-GM103540 to Michelle Digman. This work was supported in part by NIH grant RF1 AG058218 to GJB and the UC Irvine Foundation. RAS received support from the Secretaría de Educación, Ciencia, Tecnología e Innovación (SECTEI) for the elaboration of this product derived from his postdoctoral work.

## Data Available Statement

The data that support the findings of this study are available from the corresponding author upon reasonable request.

